# Model-driven promoter strength prediction based on a fine-tuned synthetic promoter library in *Escherichia coli*

**DOI:** 10.1101/2020.06.25.170365

**Authors:** Mei Zhao, Shenghu Zhou, Longtao Wu, Yu Deng

## Abstract

Promoters are one of the most critical regulatory elements controlling metabolic pathways. However, in recent years, researchers have simply perfected promoter strength, but ignored the relationship between the internal sequences and promoter strength. In this context, we constructed and characterized a mutant promoter library of P_trc_ through dozens of mutation-construction-screening-characterization engineering cycles. After excluding invalid mutation sites, we established a synthetic promoter library, which consisted of 3665 different variants, displaying an intensity range of more than two orders of magnitude. The strongest variant was 1.52-fold stronger than a 1 mM isopropyl-β-D-thiogalactoside driven P_T7_ promoter. Our synthetic promoter library exhibited superior applicability when expressing different reporters, in both plasmids and the genome. Different machine learning models were built and optimized to explore relationships between the promoter sequences and transcriptional strength. Finally, our XgBoost model exhibited optimal performance, and we utilized this approach to precisely predict the strength of artificially designed promoter sequences. Our work provides a powerful platform that enables the predictable tuning of promoters to achieve the optimal transcriptional strength.

## Introduction

The application of synthetic biology and metabolic engineering depends on essential biological regulatory elements, such as promoters^1, 2^, ribosome binding sites (RBS)^3, 4^ and terminators ^5^. These important elements make genetic circuits more tunable, inducible, responsive, and/or coordinated^6, 7, 8^. Basic levels of transcriptional regulation occur at promoters to ensure natural and synthetic circuits or metabolic pathways^9, 10^. To increase regulatory gene expression efficiency, several promoters with gradient strengths were built and modified by optimizing important genetic elements, such as −35/−10 boxes, 5’-untranslated regions and transcription factor binding sites ^11, 12, 13^. However, due to weak promoter strengths, low dynamic ranges (the highest strength/the lowest strength), limited library promoters, and inducers required, these promoters are often incapable of fine-tuning metabolic pathways. Thus, it is important to establish and characterize a comprehensive library consisting of hundreds of promoters, with continuous and broad dynamic ranges.

To overcome these limitations, Mey *et al*. constructed a synthetic promoter library with 75 variants using degenerated oligonucleotide primers, comprising a 57 bp length sequence of 20 random, 13 semi-conserved, and 24 conserved nucleotides ^14^. Although the strength of this library was 0.14- to 275-fold that of the *Escherichia coli* constitutive promoter P_LacI_, the library was small, and the strongest promoter was far lower than the commonly used P_T7_ promoter. To further extend library size, Zhou *et al*. ^15^ and Yang *et al*. ^16^ screened a hundred native promoters from *E. coli* and *Bacillus subtilis*. The transcriptional intensity ranged from 0.007%–4630% that of the P_BAD_ promoter, and 0.03–2.03-fold that of the P_43_ promoter at the transcriptional level. However, the strength of these promoters was still not comparative, or far lower than other well-studied promoters, such as P_43_ ^17^, P_Veg_ ^17^, P_T7_, P_trc_ ^18^, and P_Thl_ ^19^.

The mutation, modification, or screening of existing promoters is difficult to obtain the desired ones, thus the *de novo* design of optimized promoters from sequences is a promising approach. To do this, the relationship between the promoter sequences and intensity should be established. However, few reports have contributed to this area. For example, Jensen *et al*. ^13^ revealed a simple statistical method to explore nucleotide positions, which exerted critical effect on promoter intensity in 69 P_L-λ_ promoter variants in *E. coli*. Likewise, Mey *et al*. ^14^ and Liu *et al*. ^17^ observed similar results by analyzing a partial least squares (PLS) model in *E. coli* and *Bacillus subtilis* using 49 and 214 synthetic promoters, respectively. However, these reports suffered small data issues, single modeling, imperfect correlations and low dynamic ranges. Therefore, it is important to identify promoters with gradient strengths, broad dynamic ranges, and clear sequence profiles to explore and analyze relationships between promoter sequences and intensity, using huge data and model comparisons. Nowadays, significant advances have been made in machine and deep learning for big data analytics ^2, 20^, making promoter strength prediction achievable. In particular, Gradient Boosting Decision Tree (GDBT)^21^, AdaBoost^22^, Random Forest Regressor^23^, Xgboost^24^, and Recurrent Neural Network^25^ by one-hot coding have provided efficient mechanisms to analyze big data in designing functional promoters.

It is well known that the core promoter (−35 box, spacer, and −10 box) and their flanking regions (down and up elements) of bacterial promoters are closely related with promoter intensity ^26, 27^. To generate high strength constitutive promoters and broad dynamic range libraries, we randomized P_trc_^18^ regions by mutation-construction-screening-characterization (MCSC) engineering cycles (Fig. 1a). In doing so, P_trc_ variants we abolished its inherent limitations and derived a series of extremely high strength constitutive promoters. Based on this mutation library, we reconstructed and characterized a *de novo* synthetic promoter library which can be used as a comprehensive synthetic biology toolbox for gene expression regulation over a broad range (Fig. 1b-d). Furthermore, we investigated a direct correlation between promoter base sequences and transcription strength of the synthetic promoter library, using machine learning models (Fig. 1e). Taken together, this work not only provided the constitutive promoters, whose strengths span from low to extremely high, but also established a promoter strength prediction model which could significantly reduce promoter characterization workload.

**Fig. 1.**
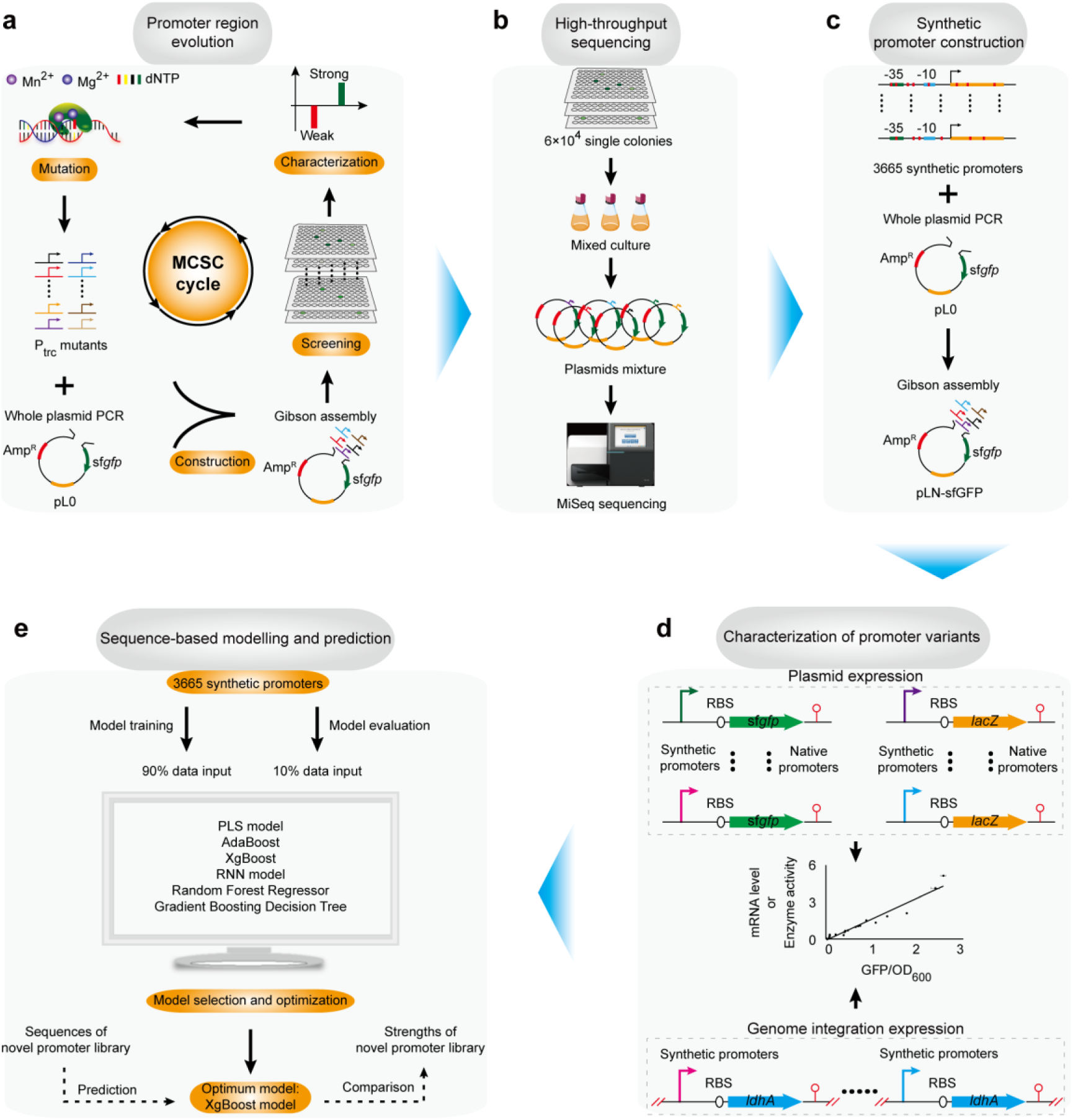
Schematic diagram of the strategies based on the promoter library to predict promoter strength. (a) Schematic illustration of the screening procedure for the promoter region evolution by MCSC engineering approach. (b) High-throughput sequencing for all mutant promoters. (c) Reconstruction of the synthetic promoter library based on MiSeq. (d) Characterization of promoter variants using different reporters. (e) Modeling and prediction based on base sequence and intensity.

## Results

### Generation of the mutant library

We chose the P_trc_ promoter (74 bp in length) ^18^ as a template from the pTrc99a plasmid to obtain a mutant constitutive promoter library by *error-prone* PCR in this work. To extend span strength of the mutation library, iterative MCSC engineering cycles (Fig. 1a) was performed to obtain both strength enhanced and reduced promoters. After each round of mutagenesis, we counted the minimum and maximum fluorescence/OD_600_ and dynamic range. The minimum fluorescence/OD_600_ was relatively stable, remaining between 85–200, while the maximum fluorescence/OD_600_ showed an obviously increasing trend, and approached the highest after 40 rounds of MCSC engineering (Fig. S1 a-b). The dynamic range reached the maximum value after 82 rounds of MCSC engineering (Fig. S1 c). After each round of MCSC engineering, we selected the mutants whose fluorescence intensities changed more than 10 times (increased or decreased) as the templates for the next round of MCSC engineering. Finally, we obtained ∼6×10^4^ constructs with an intensity range ∼509-fold difference between the strongest and weakest expression, and the strongest one had the expression levels ∼127- and 3.14-fold higher than the uninduced and induced P_trc_ (1 mM isopropyl-β-D-thiogalactoside (IPTG) induction), respectively (Fig. 2a). Therefore, this mutant library demonstrated the broad expression levels and excellent resolution.

**Fig. 2.**
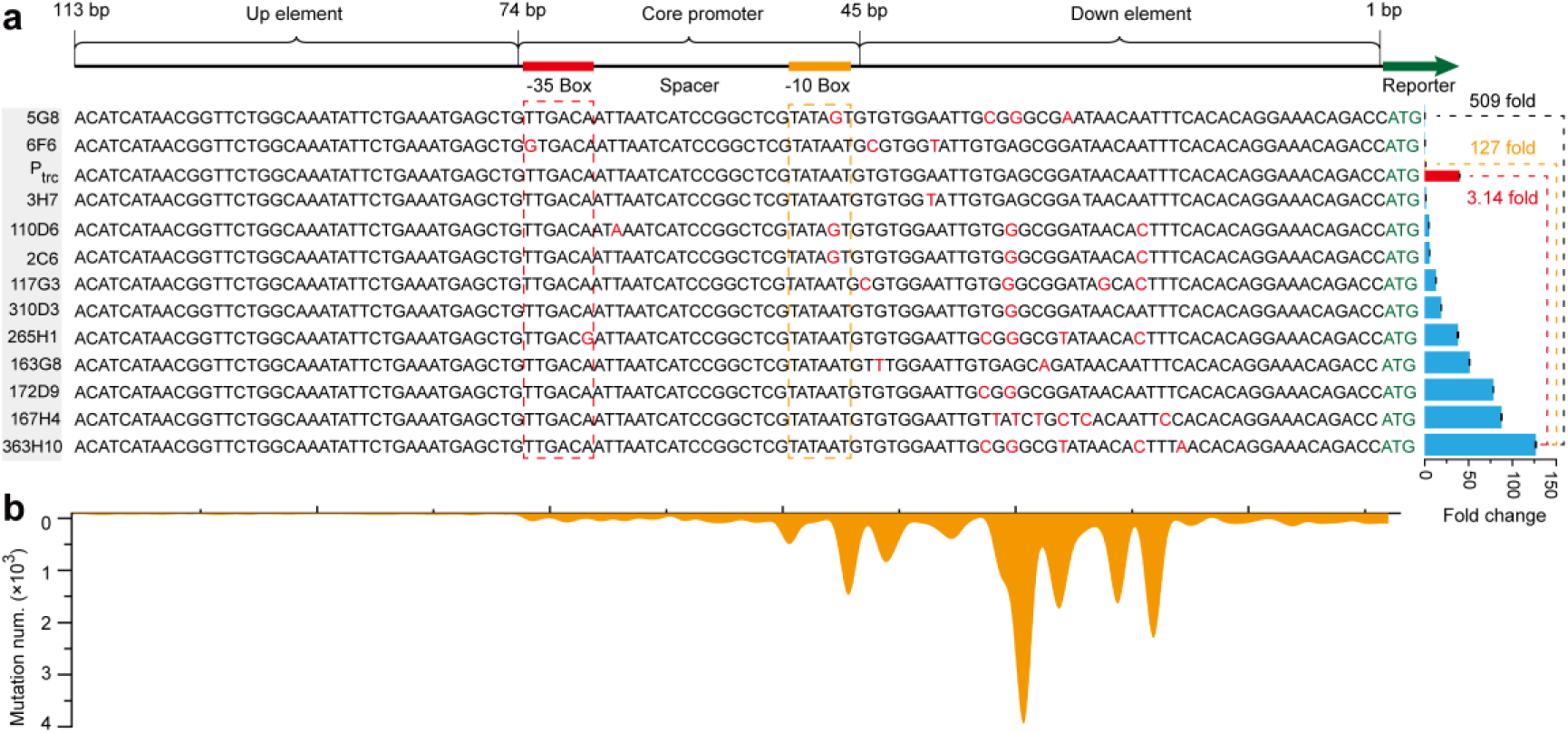
Promoter abundance distribution map. (a) Detection of fluorescence intensity and sanger sequencing for mutants during MCSC engineering. The structure of the entire promoter region included up element, core promoter, down element. 5G8 represents the strain’s position was in G8 wells of the fifth 96 plates. Other similar strains were named as described above. Red column represents 1 mM IPTG induced P_trc_ promoter. Orange column represents uninduced P_trc_ promoter. (b) Distribution of mutation positions in high-throughput sequencing.

### Sequence variances of the mutant P_trc_-derived library

After constructing the mutant P_trc_-derived library, we used the high-throughput sequencing (MiSeq) to understand the sequence variances of the mutant promoters (Fig. 1b) (SRA database # SRR11574455, https://www.ncbi.nlm.nih.gov/sra/SRR11574455). The MiSeq results show that the mutations were mainly located in the core promoter and the down element, indicating that the up element only contributes a little to the promoter strength (Fig. 2b). Furthermore, a total of 66026 different mutants were found in the library. In statistical analysis, 32.5% of the mutants contained mutations in both the core promoter and down element region, 63.06% of the mutants contained mutations only in the down element region, and 0.28% of the mutants contained mutations only in the up element region. Therefore, we focused on both the core promoter and down element to exclude the invalid mutations in the up element and reconstruct a simpler synthetic promoter library (3665 out of 66026 mutant promoters). The 3665 different P_trc_-derived mutant promoters (74-bp covering core promoter and down element) ^28^ were synthesized to form a synthetic promoter library (see Additional file 2). The 3665 different promoter sequences had 1067 mutations in −35 and/or −10 boxes, 1313 mutations in spacers, and 3581 mutations in down elements, including 9 additions, 513 deletions, and 3143 substitutions. The diverse mutations caused the diversity of promoter changes, which provides a guarantee for the evolution of promoters.

### Characterization of the synthetic promoter library

After high-throughput sequencing and reconstruction, the fluorescence intensity of all reconstructs in the synthetic promoter library were measured and analyzed. Three methods were adopted and evaluated for the synthetic promoter library. First, to standardize our synthetic promoter library, all synthetic promoters were compared with the induced P_trc_ and P_T7_ promoters in *E. coli*. To do this, P_T7_ promoter was used to replace P_Trc_ promoter on the backbone of vector pL0-sfGFP (Fig. 3a). To focus on comparing the strength of the promoter, all constructs were performed on the same backbone and introduced into *E. coli*. Expressions and comparison between fluorescent intensity of the synthetic promoters and the known promoters are shown in Fig. 3b. The maximum strength of our synthetic promoter library was ∼114-fold that of uninduced P_trc_, ∼1.52-fold that of 1 mM IPTG induced P_T7_, and ∼2.83-fold that of 1 mM IPTG induced P_trc_, with a ∼454-fold difference between the strongest and weakest expressions.

To analyze the universality of the synthetic promoter library when expressing different genes, a set of synthetic promoters (selected based on multiples of control fluorescence intensity) ^29^ were chosen and used to control the expression of β-galactosidase (*lacZ*) and lactate dehydrogenase (*ldhA*). According to the fluorescence intensity data of the synthetic promoter library, 21 promoters (PL3153, PL757, PL2776, PL862, PL3034, PL2346, PL3293, PL2983, PL1078, PL3224, PL2169, PL1958, PL1260, PL3088, PL1456, PL2666, PL3227, PL948, PL3001, PL2986, and PL3147) with fluorescence intensity multiple that of the P_trc_ (0.5–100 times) in the synthetic promoter library were selected and used to express *lacZ* instead of sf*gfp* in the plasmid (Fig. S2a and Additional file 2). After the recombinant strains were constructed, the β-galactosidase activity was measured. The same trend was observed in the β-galactosidase activities as for the sfGFP expression driven by the same promoters (Fig. S2b). The enzyme activity of *lacZ* correlated well with the fluorescence/OD_600_ (R^2^=0.94) (Fig. 3c). The specific β-galactosidase activities spanned a ∼188- and ∼60-fold range relative to the Mlac0 (MG1655ΔlacZ) and Mlac1 (pTrc-lacZ), respectively. The corresponding fluorescence intensity spanned a ∼184- and ∼71-fold range relative to the Mlac0 (MG1655ΔlacZ) and ML1 (pTrc-sfGFP), respectively. Thus, to find the most suitable promoter for expression, there is only the need to choose different folds of promoters in the synthetic library according to fluorescence intensity. Furthermore, we also found that the transcriptional (2^-ΔΔCt^) and translation level of β-galactosidase were highly correlated (Fig. 3d and Fig. S2c). The relative levels of the *lacZ* transcripts spanned a 13-fold range under the control of the selected promoters.

**Fig. 3.**
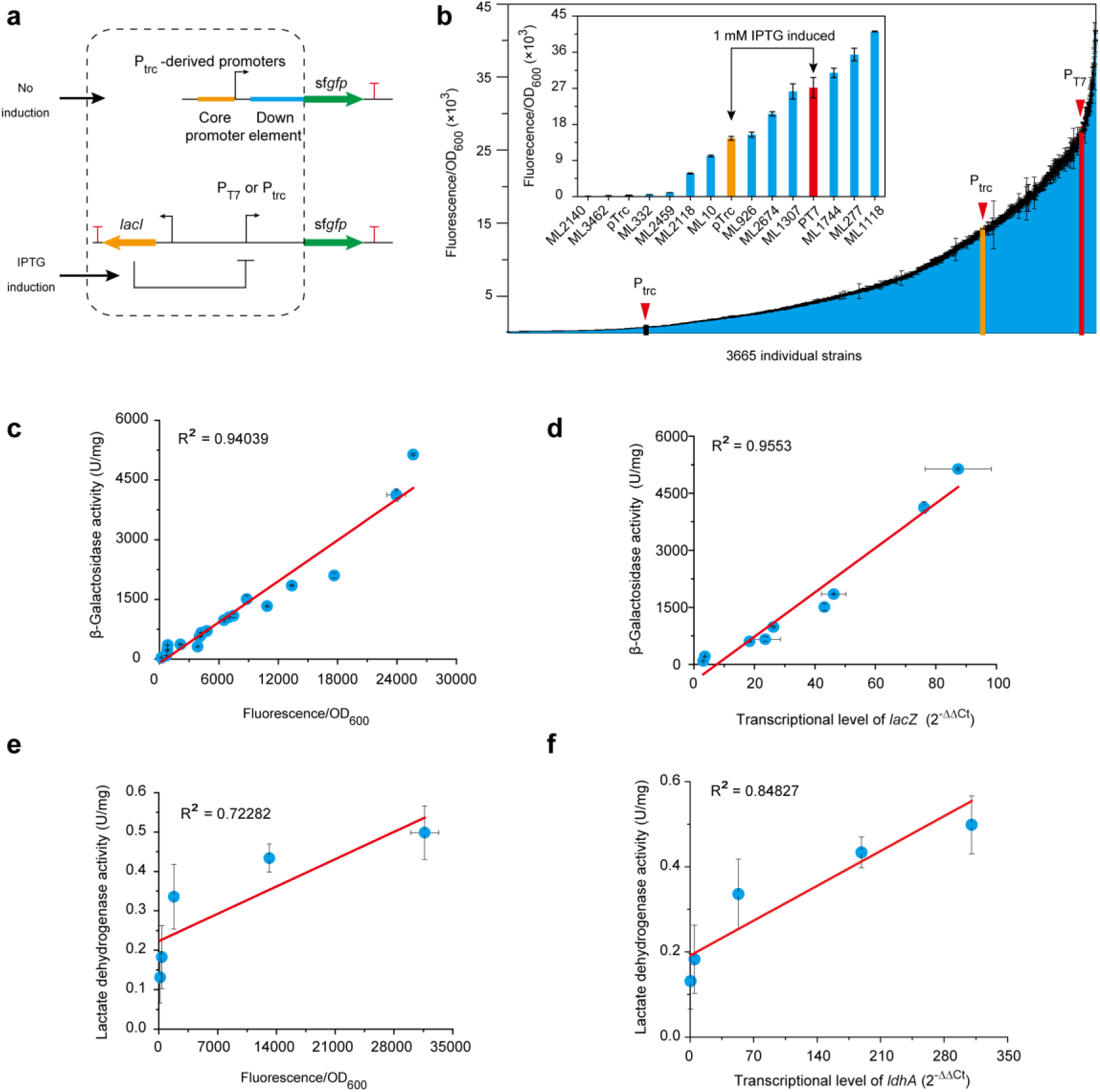
Construction and characterization of the promoter clusters using different reporters. (a) Schematic diagram of different promoters expressing the sfGFP protein. (b) Expressions and comparison of fluorescent intensity of different promoters. Data are means ± standard deviation for three independent experiments. P_T7_ and P_trc_ promoters that were induced by 1 mM IPTG were colored by red and yellow. Uninduced P_trc_ promoter was colored by black. The embedded figure represents part of the gradient strength promoters in the synthetic promoter library. (c) Relationship between β-galactosidase activity and sfGFP expression levels. (d) Cognation of the activity of promoter candidates at the transcriptional and expression level of *lacZ*. Level of changes of mRNA level (2^−ΔΔCt^) of *lacZ* measured by real-time fluorescence quantitative PCR. (e) Relationship between lactate dehydrogenase activity and sfGFP expression levels in genome. (f) Cognation of the activity of promoter candidates at the transcriptional and expression level of *ldhA*. mRNA level(2^−ΔΔCt^) of ldhA was measured by real-time fluorescence quantitative PCR. All experiments were performed three times and the error bars represent standard deviation.

To test the effect of the synthetic promoters on the genome, we modified the lactate dehydrogenase gene (*ldhA*) in *E. coli* through replacing the native *ldhA* promoter by the synthetic promoters (Fig. S3a). We deleted the wild type *ldhA* promoter in *E. coli* MG1655 forming the strain Mldh0 as control. Five different promoters, PL1409, PL908, PL2436, PL3189, and PL1993, that were 49.52%, 110.61%, 556.41%, 4086.54%, and 9826.74% the strength of P_ldhA_, respectively (Fig. S3b), were selected to substitute the P_ldhA_ in the chromosome of strain Mldh0 using CRISPR-Cas9, resulting in strains Mldh1409, Mldh2436, Mldh908, Mldh3189, and Mldh1993. The fluorescence strengths of PL2436 on the genome were almost the same as P_ldhA_, indicating the expression level of lactate dehydrogenase was similar (Fig. S3b-c). The lactate dehydrogenase activity of MG1655, Mldh1409, Mldh2436, Mldh908, Mldh3189, and Mldh1993 was 1.29, 0.96, 1.33, 2.44, 3.16, and 3.63-fold of the control (Mldh0), respectively (Fig. S3c). As shown in Fig. S3d, the transcription levels of *ldhA* (2^−ΔΔCt^) had a similar trend with fluorescence intensity and enzyme activities. Taken together, the fluorescence intensity of the same synthetic promoter candidates was tested, and the activities of lactate dehydrogenase and transcription levels of *ldhA* showed similar results (Fig. S3b-d). These results provided robust, strain-independent, gene-independent regulation.

### Machine learning-based rational design of promoters

Although a synthetic promoter library was obtained, it was extremely crucial to determine the functional relationships between promoter sequences and the strength to achieve the rational design of the desired promoter. PLS models ^11, 13, 17^ with multinomial statistics are often used for exploring nucleotide positions that have a significant effect on promoter intensity. After the promoter sequence is encoded, PLS compares the basic relationship between the two sequences. It tries to find the multi-dimensional direction of each promoter sequence to explain the multi-dimensional direction, with the largest intensity sequence variance. As such, we first tried the PLS model to predict promoter strength according to the given sequences. The data-set was randomly split into two parts: the training set, including 90% of the 3665 synthetic promoters, and the test set, including the remaining promoters ^11^. The training set was utilized to construct the PLS model. In this procedure, 74 latent variables were determined as previously described ^30^ and retained in the PLS model (R^2^=0.66) (Fig. 4a). The predicted and origin corresponded to the predicted promoter strength (log_10_(fluorescence/OD_600_)) by model and the observed promoter strength (log_10_(fluorescence/OD_600_)), respectively.

**Fig. 4.**
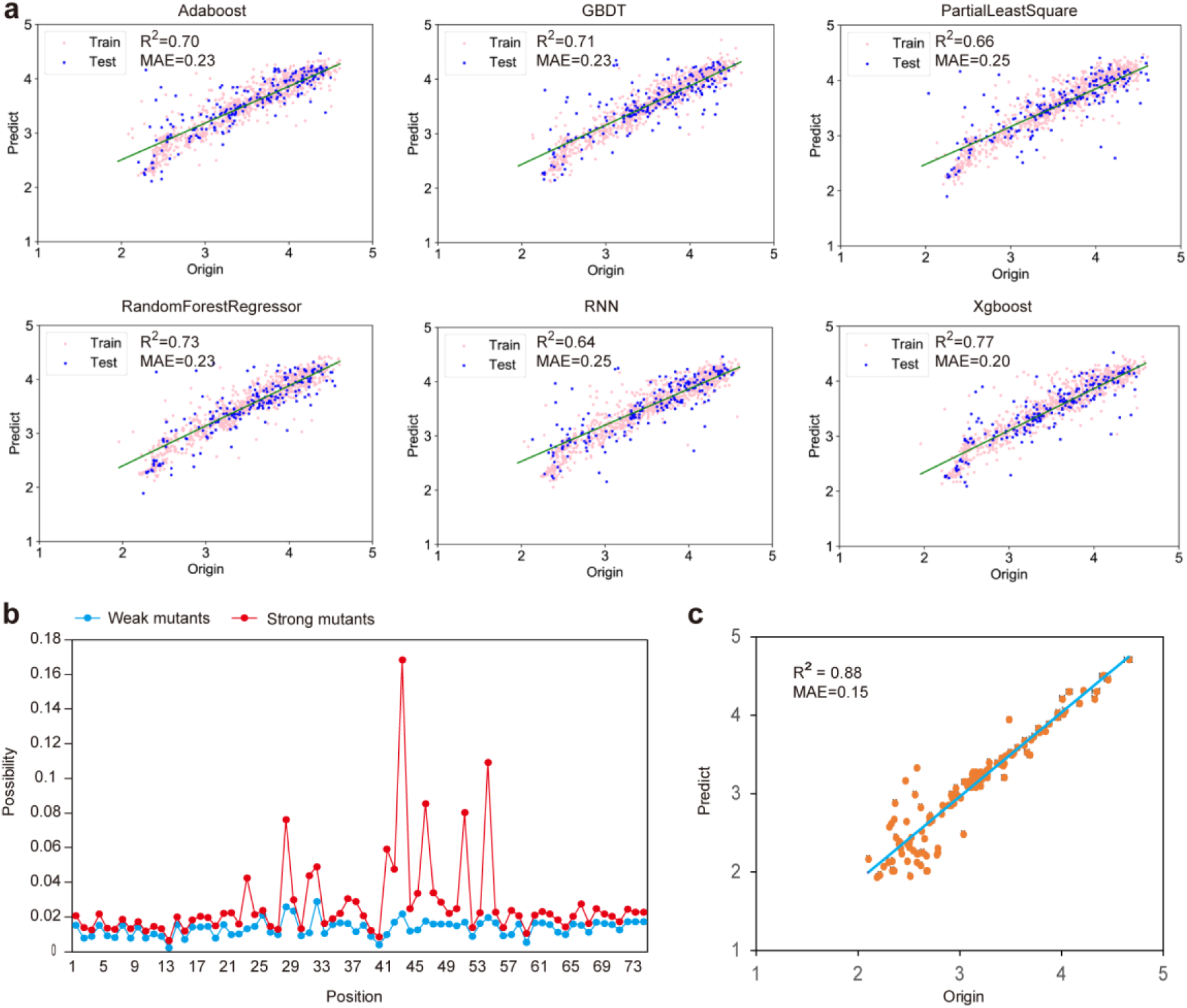
Accurate prediction of the correlation between the promoter sequence and intensity by the machine learning model. (a) Comparison and establishment of different models based on the P_trc_-derived synthetic promoter library. (b) Statistical distribution of mutations and their effects on mutant fluorescence. The red and blue curves represent strong and weak mutants, respectively. (c) The predicted promoter strength (predict, log_10_ ^(fluorescence/OD600)^) versus the observed promoter strength (origin, log_10_ ^(fluorescence/OD600)^) of the training set and the test set. The R criterion was given as the relative error sum of squares. MAE represent mean absolute error.

Due to the differences in mutant methods and huge databases, the final PLS training result was not perfect. Therefore, we started to try other machine and deep learning models, we chose the models Gradient Boosting Decision Tree (GDBT) ^21^, AdaBoost ^22^, Random Forest Regressor ^23^, XgBoost ^24^, and Recurrent Neural Network ^25^ for research using the scikit-learn Python package. All models were modeled on 90% of the data, and then the predictions of this model were tested against mean were trained to eliminate the problems of multicollinearity, and efficiently verified using 10-fold cross-validation (i.e. by training surements for the remaining 10%) with the performance measured as the mean absolute error (MAE) of relative strength (cross_val_score and RepeatedKFold from the scikit-learn Python package, https://scikit-learn.org/stable/). Finally, the XgBoost was identified as the best model with the lowest average cross-validated MAE (test MAE) (0.204) and highest R^2^ (0.77) for promoter prediction (Fig. 4a). Therefore, by applying XgBoost, an excellent relationship was found between promoter sequences and intensity. This XgBoost model can be a useful tool to rationally design a functional promoter to fine-tune gene expression. In this model, once the promoter sequence was input, the promoter strength was obtained.

### XgBoost model verification

To further verify the performances of the established XgBoost model, we rationally designed a new promoter library. To do this, the mutation possibility at each position of strong and weak promoters (the P_trc_ promoter as an interval between strong and weak ones) in the synthetic promoter library was analyzed by a statistical analysis approach ^25, 27^. The positions 25, 28, 32, 41, 43, 46, 51, and 54 were identified as the most critical bases at which mutations could significantly influence promoter strength (Fig. 4b). In this regard, we randomized these eight sites (such as A) to different bases (*i*.*e*. C, T, G, or B), leaving the other 66 sites unchanged to form a new promoter library, with a size of 390625 (5^8^ possible sequences) (Additional file 2). Predictive models of relative strength for the newly designed promoter assembly library were constructed, following a similar strategy based on one-hot encoding of features and the best model regression and cross-validation way. To accumulate enough data to train an independent model, 100 promoters (Additional file 2) were randomly selected from the new promoter assembly library, and the promoter sequence was used as an additional input feature. In addition, 27 well-studied promoters (Additional file 2) ^11, 12, 31, 32, 29, 33^ were entered into the model to test the universality. The intensity of the 127 promoters formed the test set and were predicted by the fully trained XgBoost model. The predicted intensity versus the observed intensity (fluorescence intensity) were shown in Fig. 4c. In the test set, the model generated good results. We evaluated model performance through the R criterion and the MAE (R^2^ = 0.88, MAE=0.148). These results thus indicated a satisfactory correlation of promoter strength to the sequence. Based on these validation experiments, the established XgBoost model not only held a huge numbers of experimentally verified data, but also provided a robust predictive function for the expression levels of 127 unique expression vectors in numerous conditions. Therefore, we established a relationship between promoter sequence and strength in *E. coli*, an unprecedented discovery.

## Discussion

Generally, gradient strength promoters are essential elements for pathway fine-tuning ^9, 10^. However, existing promoters suffer with low strength, narrow strength span and limited numbers. Simply engineering or screening natural promoters is not only laborious, but also difficult to identify high strength promoters. To overcome these challenges, we iteratively evolved P_trc_ promoters based on MCSC engineering cycles, and reconstructed a synthetic promoter library, after MiSeq sequencing and mutation analysis. This synthetic promoter library consisted of 3665 gradient strength promoters; the strongest promoter was 1.52-fold the strength of a 1 mM IPTG induced P_T7_ promoter. Using the synthetic promoter library as an input dataset, we built and optimized a series of promoter strength prediction models. In comparing models, the XgBoost model performed the optimally (R^2^ = 0.77, MAE = 0.205). To further verify XgBoost model reliability, we compared the predicted and actual strength of a hundred rationally designed artificial promoters (R^2^= 0.88). Taken together, we rationally designed and provided a powerful platform to enable predictable promoter tuning to transcriptional strength.

Although many advancements have been made in past decades, the strength and dynamic range of promoters were relatively low and narrow, respectively^17, 18^. In addition, promoter characterization in the literature is often performed using different genetic backgrounds and testing conditions, resulting in unquantifiable performances when applying them to the same host. Hence, several studies have screened hundreds of gradient strength constitutive promoters from *E. coli* ^15^, *B. subtilis* ^16^, *C. glutamicum* ^34^, and *S. cerevisiae* ^35^. However, the strength of these constitutive promoters is still far lower than what is required for high expression levels, especially for protein over-overexpression. Single rounds of screening from mutation libraries are difficult to obtain for extremely high strength promoters^36^. Hence, the iterative generation of high strength promoters is a promising strategy to extend promoter strength. In this regard, the P_trc_ promoter was evolved by several MCSC engineering cycles, and a series of extremely high strength promoters were screened.

We further analyzed the mutation library and found that mutation sites were mainly distributed in the core promoter (−35 box, spacer, −10 box) and down element. The core promoter is known to have a great influence on gene expression ^37, 38, 39, 40, 41, 42, 43, 44^. In-depth research found that changes in up elements ^45, 46, 47, 48^ and down elements ^49^ also had a significant contribution to gene expression, but the contribution was not as great as that of the core promoter. However, there were only a few up element changes. To facilitate the study, we only explored the core promoter and down elements of mutant promoters. We then constructed a synthetic promoter library with 3665 mutant promoter constructs, which exhibited a 454.26-fold difference between the strongest and weakest expression. The strength of the synthetic promoters was much higher than other reported promoters and it was not necessary to tandem multiple promoters to achieve a higher intensity ^17, 50^.

Although the P_trc_ promoter was generally considered as a strong inducible promoter, it worked well in the absence of an inducer ^51^. Thus, it could somehow work as a typical constitutive promoter. Originally, we searched for native constitutive promoters as the candidates for directed evolution. However, none of them was comparable with the P_Trc_ promoter in strength. In other words, the mutant native constitutive promoter might not meet our requirements of directed evolution. In this regard, we selected P_trc_ as the original promoter to establish comprehensive and constitutive promoter libraries. These libraries exhibited great stability in the expression of different reporter genes in both plasmids and the genome.

Previously, it was almost impossible to rationally predict promoter strength directly based on sequences. An ideal model was based on the predicted thermodynamics to predict the strength of the promoters. However, the thermodynamic model was too ideal to understand the promoters precisely and most of the time, the predicted promoters generated from the above models were far from the experimental results. Machine or deep learning models were independent of the “mechanisms” and thus provided a promising approach to predict promoter strength, without fully understanding mechanisms. Recently, Wang *et al*. successfully established a complicated AI model that could be used to rationally design and predict promoters^2^.

The model was based on the training of natural promoters which usually had moderate strength. Although 70.8% of promoters exhibited activity, their strength was generally low. In our study, the training of the XgBoost model was based on the high strength P_trc_ promoter library. Hence, our model predicts the high strength promoters. Furthermore, we found that the fluorescence intensity deviated from the trend line when log_10_^(sfGFP/OD600)^ lower than 2.5, which represents the low fluorescence intensity was not well worked for the model. With the development of AI, we believe that models that can precisely design desired promoters and predict the strength of a given promoter can be established in the near future.

## Methods

### Bacterial strains and cultivation

*E. coli* JM109 was used for plasmid cloning and MG1655 K12, MG1655 (DE3) were used for gene expression. Inoculates were cultured in 50 mL Luria Bertani (LB) in 250 mL shaker flasks at 200 rpm. Assay strains were inoculated (10% working volume) into M9 minimal medium ^52^, including 5 g/L D-glucose (M9G) and 0.1% amino acids^12, 31^ for the determination of fluorescence expression intensity. Gene expression was induced initially by 1 mM IPTG or no inducer ^33^. All other strains were cultured at 37°C in LB medium, supplemented with 100 mg/mL ampicillin. All strains and plasmids are listed in Table S1. All primers are listed in Table S2.

### Plasmid construction

The sfGFP ORF was amplified from the pJKR-H plasmid ^53^ with pL1-sfGFP F and pL1-sfGFP R primers and ligated into *Eco*R I/*Hin*d III sites of pTrc99a, resulting in pL1-sfGFP. The negative control and backbone pL0-sfGFP (no promoter) was constructed by whole plasmid PCR ^54^ from pL1-sfGFP, using the pL0-sfGFP F and R.

Products were digested by CIAP and ligated using DNA T4 ligase. P_T7_ promoter was amplified from pACYC-Duet-1 plasmid by primer pair of T7-F/T7-R. pT7-sfGFP plasmid was generated by whole plasmid PCR ^54^, as previously described.

Random mutagenesis, to generate novel promoters, was conducted by *error-prone* PCR ^55^ with Taq DNA polymerase in the existence of Mn^2+^, Mg^2+,^ and dNTP, using plasmid pTrc99a (Novagen, CA, USA) as the template along with the er-Trc F and er-Trc R. The primers mentioned above were used to perform 30 amplification cycles. The standard reaction conditions were as follows: 200 μl reaction volume; 10 pM each primer; 0.0625∼3 mM MnCl_2_ or 0.5∼12 mM MgCl_2_ or different ratios of 100 mM dNTP mixture; 2×Taq DNA polymerase. The cycle profile was: 1 min 94°C, 2 min 59°C, and 3 min 72°C. Then the SanPrep Column PCR Product Purification Kit (Sangon Biotech, Shanghai, China) was used to purify PCR products. The backbone was linearized by whole plasmid PCR with T0-sfGFP F and R, using plasmid pL1-sfGFP as the template. Following purification and digestion with *Dpn* I, the insert and backbone were assembled using Gibson method ^28^ and transformed into *E*.*coli* JM109. After colony PCR, these right recombinant plasmids were transferred into MG1655 and the fluorescence intensity was detected. The other constructions in this study also used the Gibson assembly method, as described above.

About ∼6×10^4^ colonies were visually screened from agar plates. A single colony from each plate was picked into M9G for fluorescence detection. After MiSeq, the reconstructed 3665 synthetic promoters were named PLN, forming the synthetic promoter library. They were transformed into MG1655 and called MLN. The promoters for predicting the XgBoost model were named pZN, transformed into MG1655, and called MZN. The synthetic promoters carrying lacZ were called pLacN, transformed into MG1655, and named MlacN. The different synthetic promoters replacing the ldhA promoter in MG1655 were called MldhN.

### MiSeq sequencing

Selected mutants were sequenced using primers Miseq F and Miseq R. A total of ∼6×10^4^ single colonies obtained by MCSC were mixed and plasmids were extracted to form a mixture sample. Samples were MiSeq sequenced by Sangon Biotech (Shanghai, China). The original MiSeq data was submitted to the SRA database, under accession number SRR11574455 (https://www.ncbi.nlm.nih.gov/sra/SRR11574455).

### Library screening using the sfGFP reporter assay

Single colonies on agar plates were inoculated into 96-well plates including 200 μL LB. After 8–12 h, the inoculums were inoculated (2% working volume) into 180 μL M9G. The cultures were grown at 37°C with 300 rpm. After 4–6 h, the fluorescence and optical density were monitored on a plate reader (Tecan) at when OD_600_ reached 0.4–0.6. A 100 μL sample was transferred to a black 96-well plate, and the sfGFP fluorescence was measured at 485 nm after excitation at 528 nm using a plate reader (Tecan). Fluorescence was measured in arbitrary units (AFU) while optical density was determined by absorbance (OD) at 600 nm. The intensity of sfGFP was characterized and calculated by sfGFP fluorescence/OD_600_. The negative controls are MG1655 K12 and ML0 (MG1655 carrying pL0-sfGFP).

### Genome manipulation

The *lacZ* and promoter *ldhA* were knocked out separately in *E. coli* MG1655 by CRISPR-Cas9 approach ^56^. Gene insertion was a similar step to the knockout procedure and the template introduced the fragment to be inserted. The template, which included the upstream 500 bp, PL908, and downstream 500 bp, was obtained from pldh908. The rest of the procedure followed the same steps as with the knockout manipulation. The synthetic promoters PL1409, PL1993, PL2436, and PL3189 were inserted in *E. coli* MG1655 using the same method. The related sgRNA was designed and shown in Supplementary Table S3.

### Analysis of transcriptional intensity

Total RNA was extracted using the Ultrapure RNA Kit (Novoprotein, Shanghai, China) and reverse transcribed using the SuperRT One-Step RT-PCR Kit (Novoprotein, Beijing, China). Real time quantitative PCR (qPCR) using a SuperRT One-Step RT-PCR Kit (Novoprotein, Beijing, China) was performed and analyzed according to the protocol ^57, 58^.

### Enzyme activity assays

Cell crude extracts were obtained and analyzed to measure enzyme activity. β-galactosidase measurements were performed as described by Miller et al. ^59, 60^. The lactate dehydrogenase assay was conducted according to previously published methods ^61, 62^. Protein concentrations were performed using the Bradford method ^63^.

### Model construction and prediction

The training set was generated by experimental parameters and calculated relative strengths. The training set contained 90% of the data, and testing predictions against measurements for the remaining 10%. Values in the training set were encoded into one-hot binary vectors. One-hot coding makes the discrete features continuous, allowing the model to optimally process data. On the other hand, through the representation of one-dimensional vectors, the purpose of expanding features is achieved, to a certain extent, the features can be sparse to prevent overfitting. Each input sequence was encoded into an eigenvector of length 74 bp with this method. The models were trained (Gradient Boosting Decision Tree (GDBT)^21^, AdaBoost^22^, Random Forest Regressor^23^, Xgboost^24^, Recurrent Neural Network^25^) to eliminate multicollinearity interference, and were cross-validated by 10-fold cross-validation with the performance indicators as the MAE of relative strength (cross_val_score and RepeatedKFold from the scikit-learn Python package). Using this data processing method for coding promoters, we established a promoter sequence predictive model based on each algorithm, to predict promoter strength using the same strategy. This code can be found at https://github.com/YuDengLAB/Predictive-the-correlation-between-promoter-base-and-intensity-through-models-comparing.

## Supporting information

Promoter information

## Data availability

All experimental data were determined in triplicate, and error bars represent the standard deviation. The original MiSeq data has been submitted to the SRA database under accession number SRR11574455 (https://www.ncbi.nlm.nih.gov/sra/SRR11574455). The code to predict the correlation between promoter base and intensity through comparing models can be found at https://github.com/YuDengLAB/Predictive-the-correlation-between-promoter-base-and-intensity-through-models-comparing.

## Acknowledgements

This work was supported by the National Key R&D Program of China (2019YFA0905502), the National Natural Science Foundation of China (21877053, 31900066), the Top-Notch Academic Programs Project of Jiangsu Higher Education Institutions (TAPP), the National First-class Discipline Program of Light Industry Technology and Engineering (LITE2018-24), the Fundamental Research Funds for the Central Universities (JUSRP51705A).

## Author contributions

Y.D. and M.Z. conceived the study; M.Z. designed and performed the experiments; S.Z. assisted the fluorescence measurement; L.W. assisted with data analysis and programs; M.Z., Y.D, and S.Z.. wrote the draft; S.Z., and M.Z. drew the figures; Y.D., S.Z., and M.Z. discussed and provided suggestions for this study. All authors reviewed, approved, and contributed to the final version of the manuscript.

## Additional information Additional file 1

Fig. S1 The promoter library profiles of each round of MCSC engineering cycles.

Fig. S2 Comparing the fluorescence, LacZ activity, and transcriptional level of the Ptrc-derived synthetic promoters.

Fig. S3 Comparing the fluorescence, LdhA activity, and transcriptional level of the Ptrc-derived synthetic promoters.

Table S1 Strains and plasmids used in this study

Table S2 Primers used in this study

## Additional file 2

3665 synthetic promoters

3665 recombinant strains

27 literature promoters

100 random assembly promoters

390625 assembly promoters

