## Supplementary Figures and Tables for "Model-driven promoter strength prediction based on a fine-tuned synthetic promoter library in *Escherichia coli*"

### This document contains:

**Figures S1-3**

**Tables S1-2**

**Supplementary References**

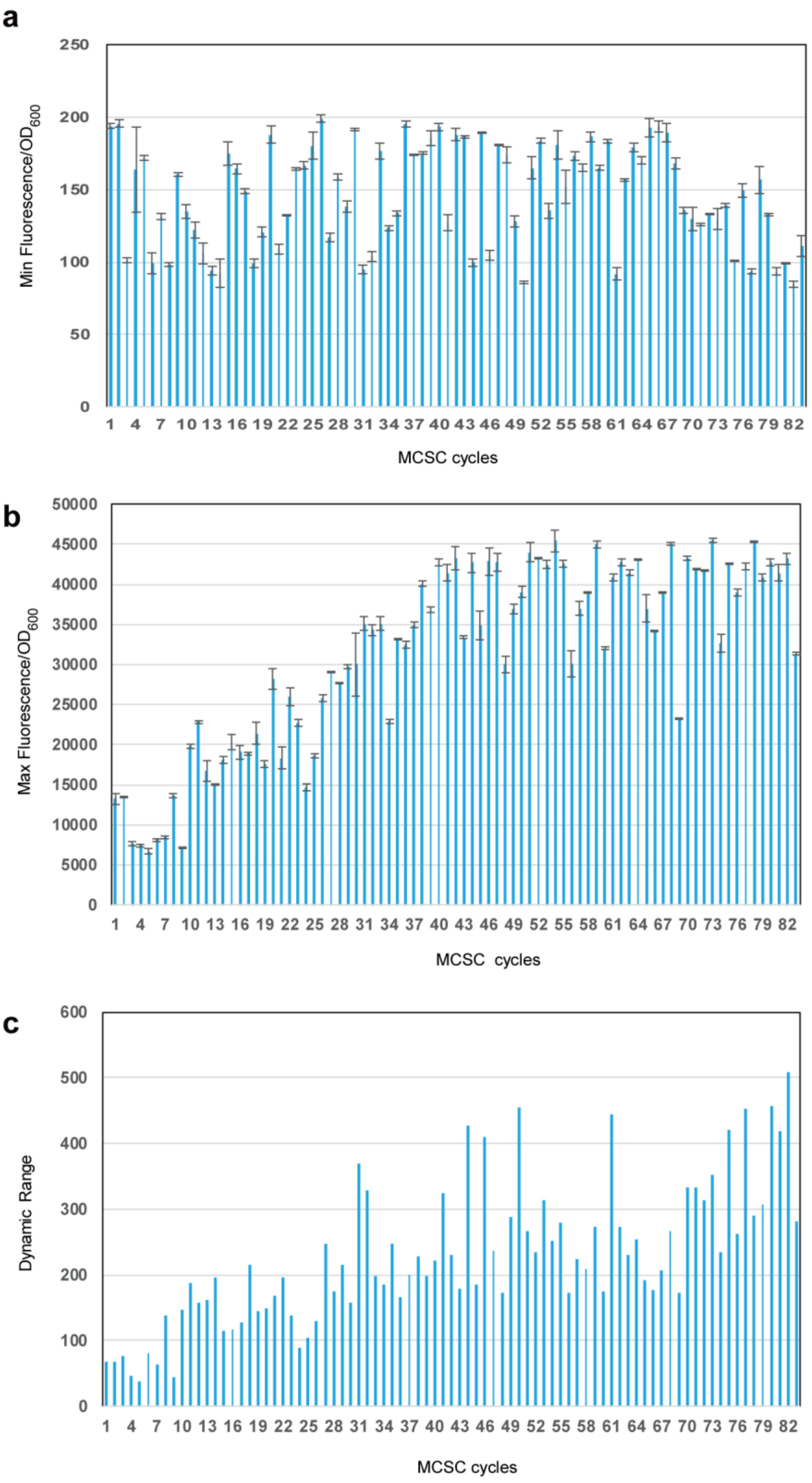


**Figure S1 The promoter library profiles of each round of MCSC engineering cycles.**

(a) The results of min fluorescence/$OD_{600}$ in every round of MCSC engineering. (b) The results of max fluorescence/$OD_{600}$ in every round of MCSC engineering. (c) The results of dynamic range in every round of MCSC engineering.

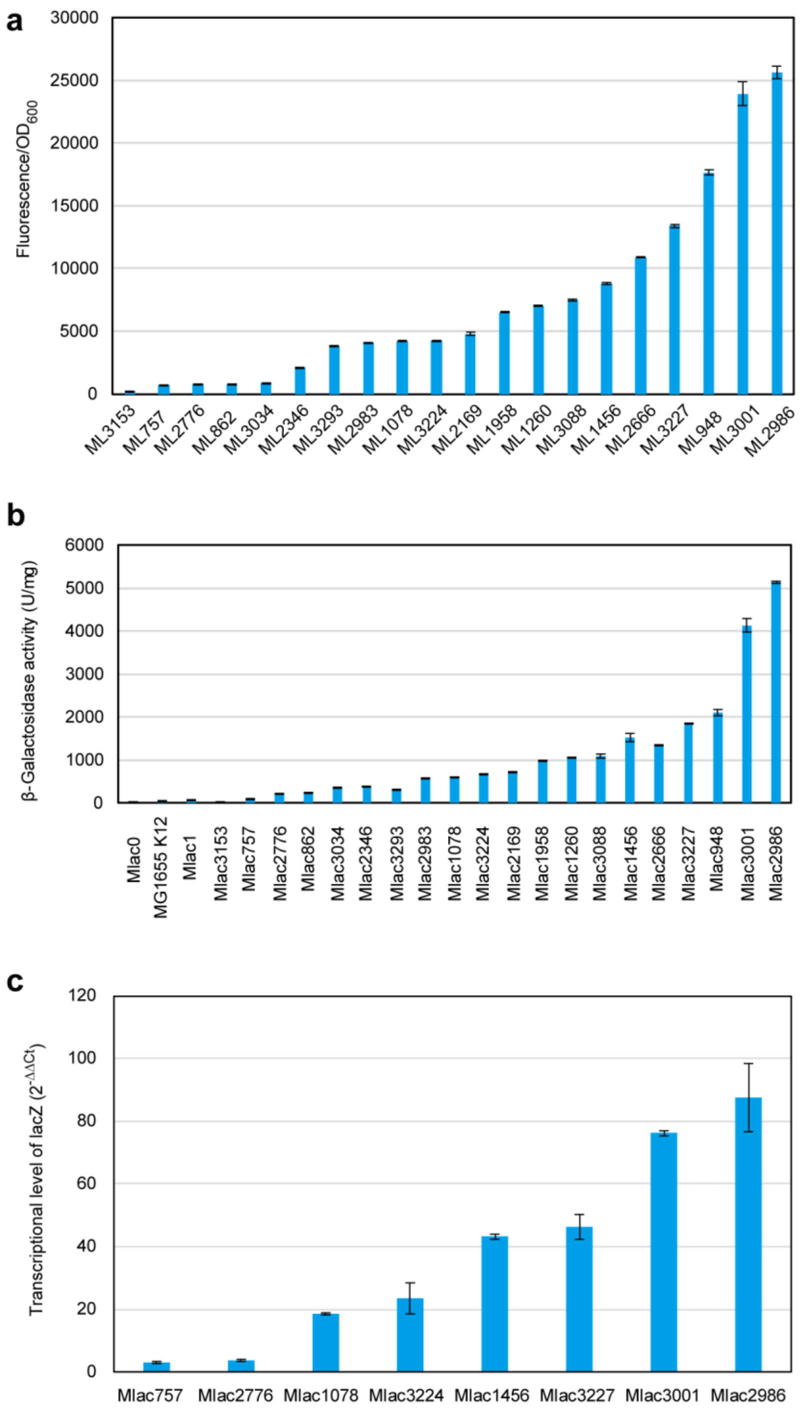


**Figure S2 Comparing the fluorescence, LacZ activity, and transcriptional level of the $P_{trc}$-derived synthetic promoters.**

(a) Analysis of fluorescence intensities of the promoter library. ML0 (MG1655 carrying pL0-sfGFP (pTrc99a harboring the sfGFP but no trc promoter)) was used as the negative control; MLN (MG1655(DE3) carrying pLN-sfGFP (pL0-sfGFP harboring Trc promoter)) was used as the positive control. (b) The activities of β-galactosidase under the control of different promoters were measured. Mlac0 was MG1655 Δ*lacZ*; Mlac1 (MG1655 Δ*lacZ* harboring pLac1 (pTrc99a harboring the *lacZ*)) was used as the positive control. (c) Changes of activity of promoter candidates at the transcriptional level of *lacZ*. Level of changes of mRNA level($2^{-\Delta\Delta Ct}$) of *lacZ* measured by real-time fluorescence quantitative PCR. All data and standard errors were derived from three independent biological replicates.

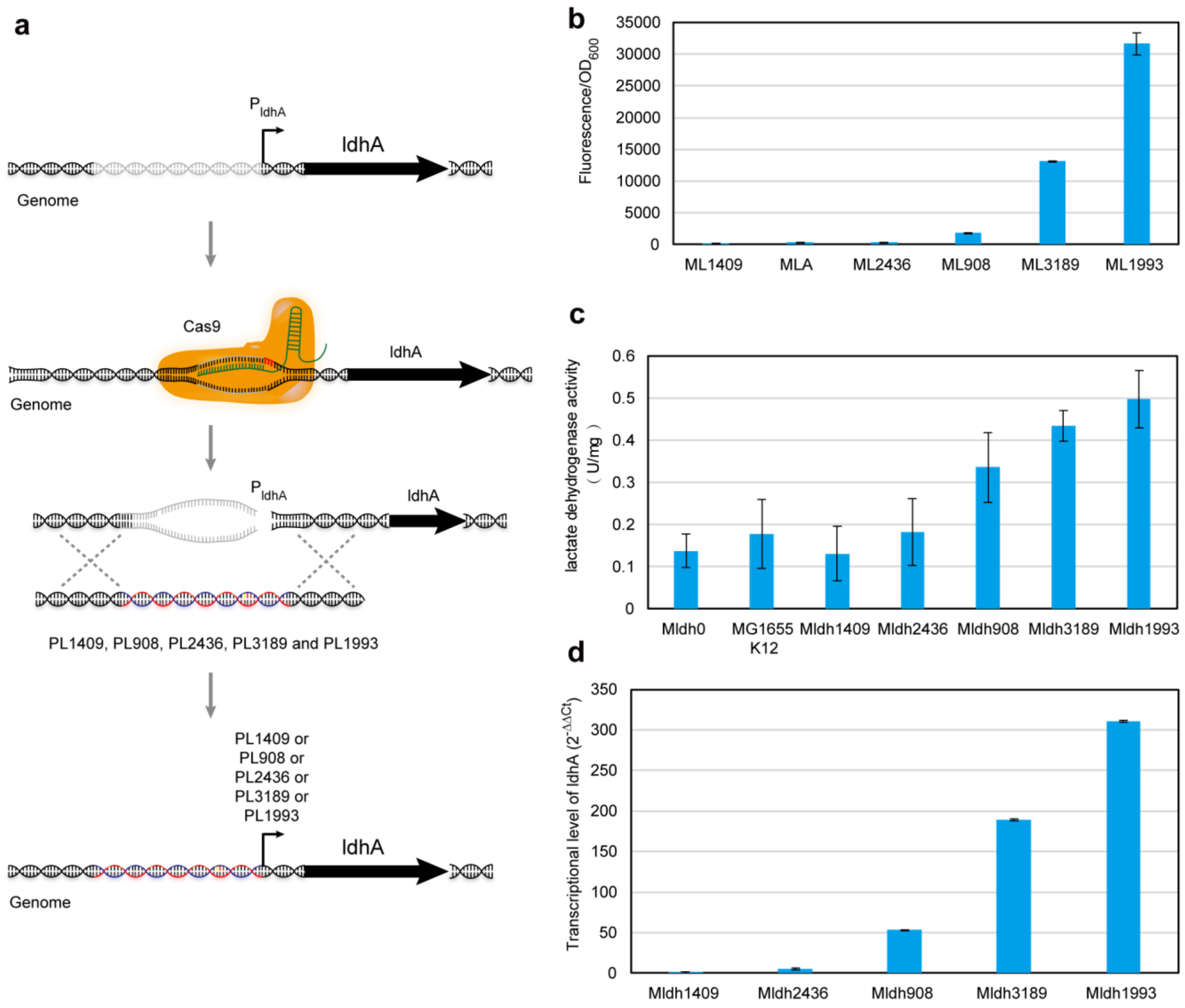


**Figure S3 Comparing the fluorescence, LdhA activity, and transcriptional level of the $P_{trc}$-derived synthetic promoters.**

(a) The promoter replacement schematic on genome. (b) Analysis of fluorescence intensities of the promoter library. ML0 (MG1655 carrying pL0-sfGFP (pTrc99a harboring the sfGFP but no trc promoter)) was used as the negative control; MLA (MG1655 carrying pLA-sfGFP (pL0-sfGFP harboring *E.coli* ldhA promoter)) was used as the positive control. (c) The activities of β-galactosidase under the control of different promoters were measured. Mldh0 (MG1655 Δ promoter ldhA) was used as the negative control; MG1655 K12 was used as the positive control. (d) Changes of activity of promoter candidates at the transcriptional level of ldhA. Level of changes of mRNA level($2^{-\Delta\Delta Ct}$) of ldhA measured by real-time fluorescence quantitative PCR. All data and standard errors were derived from three independent biological replicates.

**Table S1 Strains and plasmids used in this study**

| Strains and plasmids | Descriptions | Sources |
|---|---|---|
| **Strains** | | |
| JM109 | For plasmid construction | Novagen |
| MG1655 | For expressing genes | Novagen |
| MG1655(DE3) | For expressing genes | Novagen |
| ML0 | MG1655 carrying pL0-sfGFP | This study |
| ML1 | MG1655 carrying pL1-sfGFP | This study |
| ML2 | MG1655 carrying pL2-sfGFP | This study |
| ML3 | MG1655 carrying pL3-sfGFP | This study |
| ML4 | MG1655 carrying pL4-sfGFP | This study |
| ML5 | MG1655 carrying pL5-sfGFP | This study |
| ML6 | MG1655 carrying pL6-sfGFP | This study |
| … | … | … |
| ML3665 | MG1655 carrying pL3665-sfGFP | This study |
| MT7 | MG1655(DE3) carrying pT7-sfGFP, $Amp^R$ | This study |
| MZ1 | MG1655 carrying pZ1-sfGFP, $Amp^R$ | This study |
| MZ2 | MG1655 carrying pZ2-sfGFP, $Amp^R$ | This study |
| MZ3 | MG1655 carrying pZ3-sfGFP, $Amp^R$ | This study |
| MZ4 | MG1655 carrying pZ4-sfGFP, $Amp^R$ | This study |
| … | … | … |
| MZ127 | MG1655 carrying pZ127-sfGFP, $Amp^R$ | This study |
| Mlac0 | MG1655 $\Delta$ lacZ | This study |
| Mlac1 | MG1655 $\Delta$ lacZ harboring pLac1, $Amp^R$ | This study |

| | | |
|---|---|---|
| Mlac757 | MG1655 Δ lacZ harboring pLac757, $Amp^R$ | This study |
| Mlac862 | MG1655 Δ lacZ harboring pLac862, $Amp^R$ | This study |
| Mlac948 | MG1655 Δ lacZ harboring pLac948, $Amp^R$ | This study |
| Mlac1073 | MG1655 Δ lacZ harboring pLac1073, $Amp^R$ | This study |
| Mlac1078 | MG1655 Δ lacZ harboring pLac1078, $Amp^R$ | This study |
| Mlac1260 | MG1655 Δ lacZ harboring pLac1260, $Amp^R$ | This study |
| Mlac1456 | MG1655 Δ lacZ harboring pLac1456, $Amp^R$ | This study |
| Mlac1958 | MG1655 Δ lacZ harboring pLac1958, $Amp^R$ | This study |
| Mlac2169 | MG1655 Δ lacZ harboring pLac2169, $Amp^R$ | This study |
| Mlac2346 | MG1655 Δ lacZ harboring pLac2346, $Amp^R$ | This study |
| Mlac2666 | MG1655 Δ lacZ harboring pLac2666, $Amp^R$ | This study |
| Mlac2776 | MG1655 Δ lacZ harboring pLac2776, $Amp^R$ | This study |
| MLac2983 | MG1655 Δ lacZ harboring pLac2983, $Amp^R$ | This study |
| Mlac2986 | MG1655 Δ lacZ harboring pLac2986, $Amp^R$ | This study |
| Mlac3001 | MG1655 Δ lacZ harboring pLac3001, $Amp^R$ | This study |
| Mlac3034 | MG1655 Δ lacZ harboring pLac3034, $Amp^R$ | This study |
| Mlac3088 | MG1655 Δ lacZ harboring pLac3088, $Amp^R$ | This study |
| Mlac3147 | MG1655 Δ lacZ harboring pLac3147, $Amp^R$ | This study |
| Mlac3153 | MG1655 Δ lacZ harboring pLac3153, $Amp^R$ | This study |
| Mlac3224 | MG1655 Δ lacZ harboring pLac3224, $Amp^R$ | This study |
| Mlac3227 | MG1655 Δ lacZ harboring pLac3227, $Amp^R$ | This study |
| Mlac3293 | MG1655 Δ lacZ harboring pLac3293, $Amp^R$ | This study |
| MLA | MG1655 carrying pLA-sfGFP | This study |
| Mldh0 | MG1655 Δ promoter ldhA | This study |

| | | |
|---|---|---|
| Mldh1409 | MG1655 Δ promoter ldhA :: synthetic promoter 1409 | This study |
| Mldh908 | MG1655 Δ promoter ldhA :: synthetic promoter 908 | This study |
| Mldh1845 | MG1655 Δ promoter ldhA :: synthetic promoter 2436 | This study |
| Mldh3189 | MG1655 Δ promoter ldhA :: synthetic promoter 3189 | This study |
| Mldh1993 | MG1655 Δ promoter ldhA :: synthetic promoter 1993 | This study |
| **Plasmids** | | |
| pTrc99a | pBR322 ori, trc, $Amp^R$ | Novagen |
| pJKR-H | pUC ori, beta lactamase antibiotic resistance, High copy | (1) |
| pL0-sfGFP | pTrc99a harboring the sfGFP from pJKR-H, no trc promoter, $Amp^R$ | This study |
| pL1-sfGFP | pL0-sfGFP harboring synthetic promoter PL1(Trc), $Amp^R$ | This study |
| pL2-sfGFP | pL0-sfGFP harboring synthetic promoter PL2, $Amp^R$ | This study |
| pL3-sfGFP | pL0-sfGFP harboring synthetic promoter PL3, $Amp^R$ | This study |
| pL4-sfGFP | pL0-sfGFP harboring synthetic promoter PL4, $Amp^R$ | This study |
| pL5-sfGFP | pL0-sfGFP harboring synthetic promoter PL5, $Amp^R$ | This study |
| pL6-sfGFP | pL0-sfGFP harboring synthetic promoter PL6, $Amp^R$ | This study |
| … | … | …. |
| pL3665-sfGFP | pL0-sfGFP harboring synthetic promoter PL3665, $Amp^R$ | This study |
| pT7-sfGFP | pL0-sfGFP harboring T7 promoter, $Amp^R$ | This study |
| pZ1-sfGFP | pL0-sfGFP harboring PZ1promoter, $Amp^R$ | This study |
| pZ2-sfGFP | pL0-sfGFP harboring PZ2 promoter, $Amp^R$ | This study |
| pZ3-sfGFP | pL0-sfGFP harboring PZ3 promoter, $Amp^R$ | This study |
| pZ4-sfGFP | pL0-sfGFP harboring PZ4 promoter, $Amp^R$ | This study |
| … | … | … |

| | | |
|---|---|---|
| pZ127-sfGFP | pL0-sfGFP harboring PZ127 promoter, $Amp^R$ | This study |
| pRSFDuet-1 | RSF ori, lacI, T7 lac, $Kan^R$ | Novagen(2) |
| placZ | pRSFDuet-1 harboring the *lacZ* gene *upstream* and downstream 500bp from *E. coli*, $Kan^R$ | This study |
| pLac1 | pTrc99a harboring the lacZ from *E. coli* genome, $Amp^R$ | This study |
| pLac757 | pLac1-lacZ harboring mutant trc promoter 757, $Amp^R$ | This study |
| pLac862 | pLac1-lacZ harboring mutant trc promoter 862, $Amp^R$ | This study |
| pLac948 | pLac1-lacZ harboring mutant trc promoter 948, $Amp^R$ | This study |
| pLac1073 | pLac1-lacZ harboring mutant trc promoter 1073, $Amp^R$ | This study |
| pLac1078 | pLac1-lacZ harboring mutant trc promoter 1078, $Amp^R$ | This study |
| pLac1260 | pLac1-lacZ harboring mutant trc promoter 1260, $Amp^R$ | This study |
| pLac1456 | pLac1-lacZ harboring mutant trc promoter 1456, $Amp^R$ | This study |
| pLac1958 | pLac1-lacZ harboring mutant trc promoter 1958, $Amp^R$ | This study |
| pLac2169 | pLac1-lacZ harboring mutant trc promoter 2169, $Amp^R$ | This study |
| pLac2346 | pLac1-lacZ harboring mutant trc promoter 2346, $Amp^R$ | This study |
| pLac2666 | pLac1-lacZ harboring mutant trc promoter 2666, $Amp^R$ | This study |
| pLac2776 | pLac1-lacZ harboring mutant trc promoter 2776, $Amp^R$ | This study |
| pLac2983 | pLac1-lacZ harboring mutant trc promoter 2983, $Amp^R$ | This study |
| pLac2986 | pLac1-lacZ harboring mutant trc promoter 2986, $Amp^R$ | This study |
| pLac3001 | pLac1-lacZ harboring mutant trc promoter 3001, $Amp^R$ | This study |
| pLac3034 | pLac1-lacZ harboring mutant trc promoter 3034, $Amp^R$ | This study |
| pLac3088 | pLac1-lacZ harboring mutant trc promoter 3088, $Amp^R$ | This study |
| pLac3147 | pLac1-lacZ harboring mutant trc promoter 3147, $Amp^R$ | This study |

| | | |
|---|---|---|
| pLac3153 | pLac1-lacZ harboring mutant trc promoter 3153, Amp$^R$ | This study |
| pLac3224 | pLac1-lacZ harboring mutant trc promoter 3224, Amp$^R$ | This study |
| pLac3227 | pLac1-lacZ harboring mutant trc promoter 3227, Amp$^R$ | This study |
| pLac3293 | pLac1-lacZ harboring mutant trc promoter 3293, Amp$^R$ | This study |
| pLA-sfGFP | pL0-sfGFP harboring *E. coli* ldhA promoter, Amp$^R$ | This study |
| pldhA | pRSFDuet-1 harboring the *ldhA* gene promoter *upstream* and downstream 500bp from *E. coli*, Kan$^R$ | This study |
| pldh908 | pldhA harboring mutant promoter 908 between the *ldhA* gene promoter *upstream* and downstream 500bp, Kan$^R$ | This study |
| pldh1409 | pldhA harboring mutant promoter 1409 between the *ldhA* gene promoter *upstream* and downstream 500bp, Kan$^R$ | This study |
| pldh1993 | pldhA harboring mutant promoter 1993 between the *ldhA* gene promoter *upstream* and downstream 500bp, Kan$^R$ | This study |
| pldh2436 | pldhA harboring mutant promoter 2436 between the *ldhA* gene promoter *upstream* and downstream 500bp, Kan$^R$ | This study |
| pldh3189 | pldhA harboring mutant promoter 3189 between the *ldhA* gene promoter *upstream* and downstream 500bp, Kan$^R$ | This study |
| pCas | paraB-gam-bet-exo, bla (kan$^R$), kanR-repA101 (ts), $\lambda$-Red, the sgRNA with a lacI$^q$-Ptrc promoter guiding the pMB1 replication of pTarget | Addgene(3) |
| pTarget | the sgRNA sequence, a targeting N20 sequence, the pMB1 replicon, Spe$^R$ | Addgene(3) |

**Table S2 Primers used in this study**

| Primer | Sequence (5' to 3') |
|---|---|
| pL0-sfGFP F | ATGGAATTCATGCGTATAGGTGAAGAACTGTTCACCGGTGTTGTTC |
| pL0-sfGFP R | CAGCTCATTTCAGAATATTTGCCAGAACCG |
| pL1-sfGFP F | CAGACCATGGAATTCATGCGTATAGGTGAAGAAC |
| pL1-sfGFP R | CAAAACAGCCAAGCTTCTTTGTACAGTTCGTCCA |
| T7-F | ATATTCTGAAATGAGCTGTAATACGACTCACTATAGGGGAATTGT |
| T7-R | TATACGCATGAATTCCATGGTATATCTCCTTATTAAAGTTAAACAAAATT |
| er-Trc F | ACATCATAACGGTTCTGGCAAATATTCTGAAATGAGCTGTTGACAATTA ATCATCCGGC |
| er-Trc R | ACCGGTGAACAGTTCTTCACCTATACGCATGAATTCCATGGTCTGTTTCC TGTGTGAAA |
| pT7-sfGFP F | TAATACGACTCACTATAATGGAATTCATGCGTATAGGTGAAGAACTGTT CACCGGTG |
| pT7-sfGFP R | CAGCTCATTTCAGAATATTTGCCAGAACCG |
| Miseq F | CCTACACGACGCTCTTCCGATCTCGCCCACCGGCAGCCATCGGAAGCTG TGGTATGGCTGT |
| Miseq R | GGAGTTCCTTGGCACCCGAGAATTCCACTAGTCGGCCACGGAACCGGCA GTTTACCGGTGG |
| veri-pTrc F | AGGCAGCCATCGGAAGCTGTGGTA |
| veri-pTrc R | GAGACCCCACACTACCATCGGCG |
| veri-T0 F | GAAATGAGCTGATGGAATTCATGCGTATAGG |
| veri-lacZ | GCCCGAGTTTGTCAGAAAGCCCGTCTTCAACGTTGTGACGGTAATACGA |
| Rveri | CTCACTATAGAAATGAGCTGATGGAATTCATGCGTATAGG |

| | |
|---|---|
| Rveri-T7<br>Fveri-T0 F<br>Seq2FplacZ<br>down FplacZ<br>up Rveri<br>Rveri-T0<br>Fveri-ssrS F | AGGCAGCCATCGGAAGCTGTGGTAGAATTGTGAGCGGATAACAATTTCA<br>CACAGGAAACAGCTTAATAACCGGGCAGGCCATGAGCTGTTTCCTGTGT<br>GAAATTGTTATCCGCTCCCGTCTTCAACGTTGTGACGGGAAATGAGCTG<br>ATGGAATTCATGCGTATAGGCGGTTGCAAGATCTGAAAGAAC |
| Seq2RplacZ<br>down RplacZ<br>down FplacZ<br>up Fveri-tet<br>Fveri-T0 F | CGGGTAACGAGCGAAGCACTGAACTTTCTGTTCGACTTAAGCATTATGC<br>GGCCGCAAGCTTCCAACACAGCCAAACATCCGGAATTGTGAGCGGATA<br>ACAATTTCACACAGGAAACAGCTTAATAACCGGGCAGGCCATGCAGCC<br>ATCACCATCATCACCACAGCCAGGATCCGAATTCGGCATCGTTCCCACT<br>GCGATAGTGATAGAGATTGACATCCCGAAATGAGCTGATGGAATTCATG<br>CGTATAGG |
| q16s | |
| Fveri-lacZ<br>FplacZ down<br>RplacZ up<br>Rveri-T7 | TCTTGACATCCACAGAACTTACTTTATGCTTCCGGCTCGTACTTTCTGTTC<br>GACTTAAGCATTATGCGGCCGCAAGCTTCCAACACAGCCAAACATCCGA<br>GCTGTTTCCTGTGTGAAATTGTTATCCGCTCTAATACGACTCACTATAAG<br>TGATAGAGATTGACATCCC |
| Fveri-tet F<br>q16s<br>Rveri-lacZ<br>Rveri-lacZ<br>FplacZ down<br>Fveri | TAACCCAACATTTCACAACAGCCCGAGTTTGTCAGAAAGCACTTTATGC<br>TTCCGGCTCGTGAATTGTGAGCGGATAACAATTTCACACAGGAAACAGC<br>TTAATAACCGGGCAGGCCATGCCGTCTTCAACGTTGTGACGGTAATACG<br>ACTCACTATA |

| | |
|---|---|
| Rveri-T7 F | |
| qlacZ | TCAACATCAGCCGCTACAGTCAACGGCAGCCATCGGAAGCTGTGGTATG |
| FSeqFveri-la | GCTGTGCCCGAGTTTGTCAGAAAGCACTTTCTGTTCGACTTAAGCATTAT |
| cZ RplacZ | GCGGCCGCAAGCTTCCAACACAGCCAAACATCCGCAGCCATCACCATCA |
| down RplacZ | TCACCACAGCCAGGATCCGAATTCGGCATCGTTCCCACTGCGATCCGTC |
| up Fveri R | TTCAACGTTGTGACGG |
| qlacZ | |
| | ATTCAGCCATGTGCCTTCTTCCGTCGGCCACGGAACCGGCAGTTTACCG |
| RSeqRSeqFv | |
| | GTGGGGCAGCCATCGGAAGCTGTGGTATGGCTGTACTTTATGCTTCCGG |
| eri-lacZ | |
| | CTCGTAGCTGTTTCCTGTGTGAAATTGTTATCCGCTCCAGCCATCACCAT |
| FplacZ up | |
| | CATCACCACAGCCAGGATCCGAATTCGGCATCGTTCCCACTGCGAT |
| RplacZ up F | |
| qsfGFP | |
| | GGTGAAGGTGACGCTACCAACGAGGCAGCCATCGGAAGCTGTGGTATC |
| FSeq2FSeqR | |
| | GGCCACGGAACCGGCAGTTTACCGGTGGGCCCGAGTTTGTCAGAAAGCG |
| veri-lacZ | |
| | AATTGTGAGCGGATAACAATTTCACACAGGAAACAGCTTAATAACCGGG |
| RplacZ down | |
| | CAGGCCATGAGCTGTTTCCTGTGTGAAATTGTTATCCGCTC |
| FplacZ up R | |
| qsfGFP | GCGAAGCACTGAACACCGTAGGCGGGTAACGAGCGAAGCACTGAAGGC |
| RSeq2RSeq2 | AGCCATCGGAAGCTGTGGTAGGCAGCCATCGGAAGCTGTGGTATGGCTG |
| FSeqFplacZ | TACTTTCTGTTCGACTTAAGCATTATGCGGCCGCAAGCTTCCAACACAGC |
| down RplacZ | CAAACATCCGGAATTGTGAGCGGATAACAATTTCACACAGGAAACAGCT |
| down F | TAATAACCGGGCAGGCCATG |
| pLA Fq16s | GAAGGTTGCGCCTACACTAATCTTGACATCCACAGAACTTCGGGTAACG |
| FSeq2RSeqR | AGCGAAGCACTGATCGGCCACGGAACCGGCAGTTTACCGGTGGACTTTA |
| veri-lacZ | TGCTTCCGGCTCGTACTTTCTGTTCGACTTAAGCATTATGCGGCCGCAAG |

| | |
|---|---|
| FplacZ down | CTTCCAACACAGCCAAACATCCG |
| R | |
| pLA Rq16s | |
| Rq16s | AAGACTTTCTCCAGTGATGTTGTAACCCAACATTTCACAACATCTTGACA |
| FSeq2Fveri-l | TCCACAGAACTTAGGCAGCCATCGGAAGCTGTGGTAGCCCGAGTTTGTC |
| acZ | AGAAAGCACTTTATGCTTCCGGCTCGT |
| Rveri-lacZ F | |
| pLA-sfGFP | |
| | ACATCATAACGGTTCTGGCAAATATTCTGAAATGAGCTGGAAGGTTGCG |
| FqlacZ | |
| | CCTACACTAATCAACATCAGCCGCTACAGTCAACTAACCCAACATTTCA |
| Fq16s | |
| | CAACACGGGTAACGAGCGAAGCACTGAGGCAGCCATCGGAAGCTGTGG |
| RSeq2RSeqF | |
| | TATGGCTGTGCCCGAGTTTGTCAGAAAGC |
| veri-lacZ R | |
| pLA-sfGFP | |
| | AACACCGGTGAACAGTTCTTCACCTATACGCATGAATTCAAGACTTTCTC |
| RqlacZ | |
| | CAGTGATGTATTCAGCCATGTGCCTTCTTCCGTCAACATCAGCCGCTACA |
| RqlacZ | |
| | GTCAACTCTTGACATCCACAGAACTTTCGGCCACGGAACCGGCAGTTTA |
| Fq16s | |
| | CCGGTGGGGCAGCCATCGGAAGCTGTGGTATGGCTGT |
| FSeqRSeqF | |
| pldhA up | |
| | CAGCCATCACCATCATCACCACAGCCAGGATCCGAATTCCCCTGCCATT |
| FqsfGFP | |
| | CCTGCCAGGGGGTGAAGGTGACGCTACCAACGATTCAGCCATGTGCCTT |
| FqlacZ | |
| | CTTCCGTAACCCAACATTTCACAACAAGGCAGCCATCGGAAGCTGTGGT |
| Rq16s | |
| | ATCGGCCACGGAACCGGCAGTTTACCGGTGG |
| RSeq2FSeqR | |
| pldhA up | CTTGTCGTACTGTTTTGTGCTATAAACGGCGAGTTTCATAACTGAACGGT |
| RqsfGFP | TAAACATGCGCGAAGCACTGAACACCGTAGGGGTGAAGGTGACGCTAC |

| | |
|---|---|
| RqsfGFP | CAACGTCAACATCAGCCGCTACAGTCAACCGGGTAACGAGCGAAGCAC |
| FqlacZ | TGAAGGCAGCCATCGGAAGCTGTGGTA |
| FSeq2RSeq2 | |
| F | |
| pldhA down | |
| FpLA | |
| | ATGAAACTCGCCGTTTATAGCACGAAGGTTGCGCCTACACTAAGCGAAG |
| FqsfGFP | |
| | CACTGAACACCGTAGGATTCAGCCATGTGCCTTCTTCCGTCTTGACATCC |
| RqlacZ | |
| | ACAGAACTTCGGGTAACGAGCGAAGCACTGA |
| Rq16s | |
| FSeq2R | |
| pldhA down | |
| | ACTTTCTGTTCGACTTAAGCATTATGCGGCCGCAAGCTTCCAAAACCTTT |
| RpLA RpLA | |
| | CAGAATGCGAAGACTTTCTCCAGTGATGTTGGAAGGTTGCGCCTACACT |
| FqsfGFP | |
| | AAGGTGAAGGTGACGCTACCAACGTAACCCAACATTTCACAACATCTTG |
| Fq16s Rq16s | |
| | ACATCCACAGAACTT |
| F | |
| pldh908 | |
| | TGTGGGCGGATAACACTTTCACACATGAAACAGACCATGAAACTCGCCG |
| FpLA-sfGFP | |
| | TTTATAGCACACATCATAACGGTTCTGGCAAATATTCTGAAATGAGCTG |
| FpLA | |
| | GAAGGTTGCGCCTACACTAAAAGACTTTCTCCAGTGATGTTGGCGAAGC |
| RqsfGFP | |
| | ACTGAACACCGTAGGTCAACATCAGCCGCTACAGTCAACTAACCCAACA |
| RqlacZ | |
| | TTTCACAACA |
| Fq16s R | |
| pldh908 | ATTCCACACACTATACGAGCCGGATGATTAATTGTCAAAACTGAACGGT |
| RpLA-sfGFP | TAAACATGCCAACACCGGTGAACAGTTCTTCACCTATACGCATGAATTC |
| RpLA-sfGFP | AAGACTTTCTCCAGTGATGTACATCATAACGGTTCTGGCAAATATTCTGA |

| | |
|---|---|
| FpLA | AATGAGCTGGAAGGTTGCGCCTACACTAAGAAGGTTGCGCCTACACTAA |
| FqlacZ | ATTCAGCCATGTGCCTTCTTCCGTCAACATCAGCCGCTACAGTCAAC |
| RqlacZ F | |
| pldh1409 | |
| | TGTTGTTCCGATCCTGGTTGAACTGGACGGTGACGTATGAAACTCGCCG |
| FpldhA up | |
| | TTTATAGCACCAGCCATCACCATCATCACCACAGCCAGGATCCGAATTC |
| FpLA-sfGFP | |
| | CCCTGCCATTCCTGCCAGGGAACACCGGTGAACAGTTCTTCACCTATAC |
| RpLA | |
| | GCATGAATTCAAGACTTTCTCCAGTGATGTAAGACTTTCTCCAGTGATGT |
| RqsfGFP | |
| | TGGGTGAAGGTGACGCTACCAACGATTCAGCCATGTGCCTTCTTCCG |
| FqlacZ R | |
| pldh1409 | CCGGTGAACGGTTCTTCACCTATACGCATGAATTCCATAACTGAACGGT |
| RpldhA up | TAAACATGCCCTTGTCGTACTGTTTTGTGCTATAAACGGCGAGTTTCATA |
| RpldhA up | ACTGAACGGTTAAACATGCCAGCCATCACCATCATCACCACAGCCAGGA |
| FpLA-sfGFP | TCCGAATTCCCCTGCCATTCCTGCCAGGGACATCATAACGGTTCTGGCA |
| FqsfGFP | AATATTCTGAAATGAGCTGGAAGGTTGCGCCTACACTAAGCGAAGCACT |
| RqsfGFP F | GAACACCGTAGGGGTGAAGGTGACGCTACCAACG |
| pldh1993 | |
| FpldhA | GTGAGTGGATAACAATTTCACACAGGAAACAGACGATGAAACTCGCCG |
| down | TTTATAGCACAATGAAACTCGCCGTTTATAGCACCTTGTCGTACTGTTTT |
| FpldhA up | GTGCTATAAACGGCGAGTTTCATAACTGAACGGTTAAACATGCAACACC |
| RpLA-sfGFP | GGTGAACAGTTCTTCACCTATACGCATGAATTCAAGACTTTCTCCAGTGA |
| RpLA | TGTGAAGGTTGCGCCTACACTAAGCGAAGCACTGAACACCGTAGG |
| FqsfGFP R | |
| pldh1993 | AATTCCACACATTATACGAGCCGGATGATTAATTGTCAAAACTGAACGG |
| RpldhA | TTAAACATGCACTTTCTGTTCGACTTAAGCATTATGCGGCCGCAAGCTTC |

| | |
|---|---|
| down | CAAAACCTTTCAGAATGCGATGAAACTCGCCGTTTATAGCACCAGCCAT |
| RpldhA | CACCATCATCACCACAGCCAGGATCCGAATTCCCCTGCCATTCCTGCCA |
| down | GGGAAGACTTTCTCCAGTGATGTTGGAAGGTTGCGCCTACACTAA |
| FpldhA up | |
| FpLA RpLA | |
| F | |
| pldh2436 | TGTGAGCGGATAACAATTTCACACAGGAAACAGACCATGAAACTCGCC |
| Fpldh908 | GTTTATAGCACTGTGGGCGGATAACACTTTCACACATGAAACAGACCAT |
| FpldhA | GAAACTCGCCGTTTATAGCACACTTTCTGTTCGACTTAAGCATTATGCGG |
| down | CCGCAAGCTTCCAAAACCTTTCAGAATGCGCTTGTCGTACTGTTTTGTGC |
| RpldhA up | TATAAACGGCGAGTTTCATAACTGAACGGTTAAACATGCACATCATAAC |
| RpLA-sfGFP | GGTTCTGGCAAATATTCTGAAATGAGCTGGAAGGTTGCGCCTACACTAA |
| FpLA R | AAGACTTTCTCCAGTGATGTTG |
| pldh2436 | |
| | ATTCCACACATTATACGAGCCGAATGATTAATTGTCAAAACTGAACGGT |
| Rpldh908 | |
| | TAAACATGCCATTCCACACACTATACGAGCCGGATGATTAATTGTCAAA |
| Rpldh908 | |
| | ACTGAACGGTTAAACATGCCTGTGGGCGGATAACACTTTCACACATGAA |
| FpldhA | |
| | ACAGACCATGAAACTCGCCGTTTATAGCACATGAAACTCGCCGTTTATA |
| down | |
| | GCACAACACCGGTGAACAGTTCTTCACCTATACGCATGAATTCAAGACT |
| FpLA-sfGFP | |
| | TTCTCCAGTGATGTACATCATAACGGTTCTGGCAAATATTCTGAAATGA |
| RpLA-sfGFP | |
| | GCTGGAAGGTTGCGCCTACACTAA |
| F | |
| pldh3189 | TGTAAGCGGATAACAATTTCACACAGGAAACAGACCATGAAACTCGCC |
| Fpldh1409 | GTTTATAGCACTGTTGTTCCGATCCTGGTTGAACTGGACGGTGACGTATG |
| Fpldh908 | AAACTCGCCGTTTATAGCACATTCCACACACTATACGAGCCGGATGATT |

| | |
|---|---|
| RpldhA | AATTGTCAAAACTGAACGGTTAAACATGCCACTTTCTGTTCGACTTAAG |
| down | CATTATGCGGCCGCAAGCTTCCAAAACCTTTCAGAATGCGCAGCCATCA |
| RpldhA up | CCATCATCACCACAGCCAGGATCCGAATTCCCCTGCCATTCCTGCCAGG |
| FpLA-sfGFP | GAACACCGGTGAACAGTTCTTCACCTATACGCATGAATTCAAGACTTTC |
| R | TCCAGTGATGT |
| | ATTCACACATTATACGAGCCGGATGATTAATTGTCAAAACTGAACGGTT |
| pldh3189 | AAACATGCCACCGGTGAACGGTTCTTCACCTATACGCATGAATTCCATA |
| Rpldh1409 | ACTGAACGGTTAAACATGCCTGTTGTTCCGATCCTGGTTGAACTGGACG |
| Rpldh1409 | GTGACGTATGAAACTCGCCGTTTATAGCACTGTGGGCGGATAACACTTT |
| Fpldh908 | CACACATGAAACAGACCATGAAACTCGCCGTTTATAGCACCTTGTCGTA |
| FpldhA up | CTGTTTTGTGCTATAAACGGCGAGTTTCATAACTGAACGGTTAAACATG |
| RpldhA up F | CCAGCCATCACCATCATCACCACAGCCAGGATCCGAATTCCCCTGCCAT |
| | TCCTGCCAGGG |
| veri-ldhA | |
| | CGCCATAGCTTTCAATTAAATTTGGTGAGTGGATAACAATTTCACACAG |
| Fpldh1993 | |
| | GAAACAGACGATGAAACTCGCCGTTTATAGCACACCGGTGAACGGTTCT |
| Fpldh1409 | |
| | TCACCTATACGCATGAATTCCATAACTGAACGGTTAAACATGCCATTCC |
| Rpldh908 | |
| | ACACACTATACGAGCCGGATGATTAATTGTCAAAACTGAACGGTTAAAC |
| RpldhA | |
| | ATGCCATGAAACTCGCCGTTTATAGCACCTTGTCGTACTGTTTTGTGCTA |
| down | |
| | TAAACGGCGAGTTTCATAACTGAACGGTTAAACATGC |
| FpldhA up R | |
| veri-ldhA | CTTTAATAAGGAGATATACCATGAATTCCACACATTATACGAGCCGGAT |
| Fpldh1993 | GATTAATTGTCAAAACTGAACGGTTAAACATGCGTGAGTGGATAACAAT |
| Rpldh1993 | TTCACACAGGAAACAGACGATGAAACTCGCCGTTTATAGCACATGTTGT |
| Fpldh1409 | TCCGATCCTGGTTGAACTGGACGGTGACGTATGAAACTCGCCGTTTATA |

FpldhA GCACACTTTCTGTTCGACTTAAGCATTATGCGGCCGCAAGCTTCCAAAA

down CCTTTCAGAATGCGATGAAACTCGCCGTTTATAGCAC

RpldhA

down F

veri-ldh908 GTGTGTGGAATTGTGGGCGGATAACTGTGAGCGGATAACAATTTCACAC

Fpldh2436 AGGAAACAGACCATGAAACTCGCCGTTTATAGCACAATTCCACACATTA

Fpldh1993 TACGAGCCGGATGATTAATTGTCAAAACTGAACGGTTAAACATGCCCGG

Rpldh1409 TGAACGGTTCTTCACCTATACGCATGAATTCCATAACTGAACGGTTAAA

Rpldh908 CATGCCTGTGGGCGGATAACACTTTCACACATGAAACAGACCATGAAAC

FpldhA TCGCCGTTTATAGCACACTTTCTGTTCGACTTAAGCATTATGCGGCCGCA

down R AGCTTCCAAAACCTTTCAGAATGCG

CCGTTCACCGGTGTTGTTCCGATCCATTCCACACATTATACGAGCCGAAT

veri-ldh1409

GATTAATTGTCAAAACTGAACGGTTAAACATGCCTGTGAGCGGATAACA

Fpldh2436

ATTTCACACAGGAAACAGACCATGAAACTCGCCGTTTATAGCACGTGAG

Rpldh2436

TGGATAACAATTTCACACAGGAAACAGACGATGAAACTCGCCGTTTATA

Fpldh1993

GCACAATTCCACACACTATACGAGCCGGATGATTAATTGTCAAAACTGA

Fpldh908

ACGGTTAAACATGCCTGTGGGCGGATAACACTTTCACACATGAAACAGA

Rpldh908 F

CCATGAAACTCGCCGTTTATAGCAC

veri-ldh1993 AATGTGTGGAATTGTGAGTGGATAACTGTAAGCGGATAACAATTTCACA

Fpldh3189 CAGGAAACAGACCATGAAACTCGCCGTTTATAGCACATTCCACACATTA

Fpldh2436 TACGAGCCGAATGATTAATTGTCAAAACTGAACGGTTAAACATGCCAAT

Rpldh1993 TCCACACATTATACGAGCCGGATGATTAATTGTCAAAACTGAACGGTTA

Rpldh1409 AACATGCTGTTGTTCCGATCCTGGTTGAACTGGACGGTGACGTATGAAA

Fpldh908 R CTCGCCGTTTATAGCACATTCCACACACTATACGAGCCGGATGATTAATT

GTCAAAACTGAACGGTTAAACATGCC

TGTGTGGAATTGTGAGCGGATAACATTCACACATTATACGAGCCGGATG

veri-ldh2436

ATTAATTGTCAAAACTGAACGGTTAAACATGCCATGTAAGCGGATAACA

Fpldh3189

ATTTCACACAGGAAACAGACCATGAAACTCGCCGTTTATAGCACTGTGA

Rpldh3189

GCGGATAACAATTTCACACAGGAAACAGACCATGAAACTCGCCGTTTAT

Fpldh2436

AGCACCCGGTGAACGGTTCTTCACCTATACGCATGAATTCCATAACTGA

Fpldh1409

ACGGTTAAACATGCCTGTTGTTCCGATCCTGGTTGAACTGGACGGTGAC

Rpldh1409 F

GTATGAAACTCGCCGTTTATAGCAC

veri-ldh3189 ATGTGTGAATTGTAAGCGGATAACCGCCATAGCTTTCAATTAAATTTGA

Fveri-ldhA TTCACACATTATACGAGCCGGATGATTAATTGTCAAAACTGAACGGTTA

Fpldh3189 AACATGCCAATTCCACACATTATACGAGCCGAATGATTAATTGTCAAAA

Rpldh2436 CTGAACGGTTAAACATGCCGTGAGTGGATAACAATTTCACACAGGAAAC

Rpldh1993 AGACGATGAAACTCGCCGTTTATAGCACACCGGTGAACGGTTCTTCACC

Fpldh1409 R TATACGCATGAATTCCATAACTGAACGGTTAAACATGCC

veri-ldh CTTCTTATACTTAACTAATATACTAAGACTTTAATAAGGAGATATACCAT

Rveri-ldhA GCGCCATAGCTTTCAATTAAATTTGTGTAAGCGGATAACAATTTCACAC

Fveri-ldhA AGGAAACAGACCATGAAACTCGCCGTTTATAGCACAATTCCACACATTA

Fpldh3189 TACGAGCCGGATGATTAATTGTCAAAACTGAACGGTTAAACATGCGTGA

Fpldh1993 GTGGATAACAATTTCACACAGGAAACAGACGATGAAACTCGCCGTTTAT

Rpldh1993 F AGCACA

qldhA GGCGTTCGATCCGTATCCAAGTGGTGTGTGGAATTGTGGGCGGATAACC

Fveri-ldh908 TTTAATAAGGAGATATACCATGATTCACACATTATACGAGCCGGATGAT

Fveri-ldhA TAATTGTCAAAACTGAACGGTTAAACATGCCATGTGAGCGGATAACAAT

Fpldh3189 TTCACACAGGAAACAGACCATGAAACTCGCCGTTTATAGCACAATTCCA

| | |
|---|---|
| Rpldh2436 | CACATTATACGAGCCGGATGATTAATTGTCAAAACTGAACGGTTAAACA |
| Fpldh1993 R | TGC |
| qldhA | |
| Rveri-ldh140 | AGGCGGCTTCGTTCAACAGATGCCGTTCACCGGTGTTGTTCCGATCCGTG |
| 9 | TGTGGAATTGTGGGCGGATAACCGCCATAGCTTTCAATTAAATTTGATTC |
| Fveri-ldh908 | CACACATTATACGAGCCGAATGATTAATTGTCAAAACTGAACGGTTAAA |
| Fveri-ldhA | CATGCCTGTGAGCGGATAACAATTTCACACAGGAAACAGACCATGAAA |
| Fpldh2436 | CTCGCCGTTTATAGCAC |
| Rpldh2436 F | |
| sgRNA1(ldh | |
| | TAAATGTGATTCAACATCAC |
| A)veri-ldh19 | |
| | TGGAATGTGTGGAATTGTGAGTGGATAACCCGTTCACCGGTGTTGTT |
| 93 | |
| | CCGATCCCTTTAATAAGGAGATATACCATGTGTAAGCGGATAACAATTT |
| Fveri-ldh140 | |
| | CACACAGGAAACAGACCATGAAACTCGCCGTTTATAGCACATTCCACAC |
| 9 Fveri-ldhA | |
| | ATTATACGAGCCGAATGATTAATTGTCAAAACTGAACGGTTAAACATGC |
| Fpldh3189 | |
| | C |
| Fpldh2436 R | |
| sgRNA2(ldh | |
| A)veri-ldh24 | TCAAATTTAATTGAAAGCTA |
| 36 | TGGTGTGTGGAATTGTGAGCGGATAACAATGTGTGGAATTGTGAGT |
| Fveri-ldh199 | GGATAACGTGTGTGGAATTGTGGGCGGATAACATTCACACATTATACGA |
| 3 | GCCGGATGATTAATTGTCAAAACTGAACGGTTAAACATGCCATGTAAGC |
| Fveri-ldh908 | GGATAACAATTTCACACAGGAAACAGACCATGAAACTCGCCGTTTATAG |
| Fpldh3189 | CAC |
| Rpldh3189 F | |

| | |
|---|---|
| veri-ldh3189<br>Fveri-ldh243<br>6<br>Fveri-ldh140<br>9 Fveri-ldhA<br>Fpldh3189 R | ATGTGTGAATTGTAAGCGGATAACTGTGTGGAATTGTGAGCGGATAACC<br>CGTTCACCGGTGTTGTTCCGATCCCGCCATAGCTTTCAATTAAATTTGAT<br>TCACACATTATACGAGCCGGATGATTAATTGTCAAAACTGAACGGTTAA<br>ACATGCCA |
| veri-ldh<br>Rveri-ldh318<br>9<br>Fveri-ldh199<br>3 Fveri-ldhA<br>Fveri-ldhA F<br>qldhA | CTTCTTATACTTAACTAATATACTAAGAATGTGTGAATTGTAAGCGGATA<br>ACAATGTGTGGAATTGTGAGTGGATAACCTTTAATAAGGAGATATACCA<br>TGCGCCATAGCTTTCAATTAAATTTG |
| Fveri-ldh<br>Rveri-ldh243<br>6<br>Fveri-ldh908<br>Fveri-ldhA F<br>qldhA | GGCGTTCGATCCGTATCCAAGTGCTTCTTATACTTAACTAATATACTAAG<br>ATGTGTGGAATTGTGAGCGGATAACGTGTGTGGAATTGTGGGCGGATAA<br>CCTTTAATAAGGAGATATACCATG |
| RqldhA<br>Fveri-ldh318<br>9<br>Fveri-ldh140<br>9 | AGGCGGCTTCGTTCAACAGATGGGCGTTCGATCCGTATCCAAGTGATGT<br>GTGAATTGTAAGCGGATAACCCGTTCACCGGTGTTGTTCCGATCCGTGT<br>GTGGAATTGTGGGCGGATAAC |

| | |
|---|---|
| Fveri-ldh908<br>F<br>sgRNA1(ldh<br>A)qldhA<br>Rveri-ldh<br>Rveri-ldh199<br>3<br>Fveri-ldh140<br>9 F | TAAATGTGATTCAACATCAC<br>TGGAGGCGGCTTCGTTCAACAGATGCTTCTTATACTTAACTAATATA<br>CTAAGAAATGTGTGGAATTGTGAGTGGATAACCCGTTCACCGGTGTTGT<br>TCCGATCC |
| sgRNA2(ldh<br>A)sgRNA1(l<br>dhA)qldhA<br>Fveri-ldh243<br>6<br>Fveri-ldh199<br>3 F | TCAAATTTAATTGAAAGCTA TGGTAAATGTGATTCAACATCAC<br>TGGGGCGTTCGATCCGTATCCAAGTGTGTGTGGAATTGTGAGCGGAT<br>AACAATGTGTGGAATTGTGAGTGGATAAC |
| sgRNA2(ldh<br>A)qldhA<br>Rveri-ldh318<br>9<br>Fveri-ldh243<br>6 F | TCAAATTTAATTGAAAGCTA<br>TGGAGGCGGCTTCGTTCAACAGATGATGTGTGAATTGTAAGCGGAT<br>AACTGTGTGGAATTGTGAGCGGATAAC |
| sgRNA1(ldh<br>A)veri-ldh | TAAATGTGATTCAACATCAC<br>TGGCTTCTTATACTTAACTAATATACTAAGAATGTGTGAATTGTAAG |

| | |
|---|---|
| Rveri-ldh3189 F | CGGATAAC |
| sgRNA2(ldhA)qldhA Fveri-ldh R | TCAAATTTAATTGAAAGCTA TGGGGCGTTCGATCCGTATCCAAGTGCTTCTTATACTTAACTAATAT ACTAAGA |
| qldhA RqldhA F | AGGCGGCTTCGTTCAACAGATGGGCGTTCGATCCGTATCCAAGTG |
| sgRNA1(ldhA)qldhA R | TAAATGTGATTCAACATCAC TGGAGGCGGCTTCGTTCAACAGATG |
| sgRNA2(ldhA)sgRNA1(ldhA) | TCAAATTTAATTGAAAGCTA TGGTAAATGTGATTCAACATCAC TGG |
| sgRNA2(ldhA) | TCAAATTTAATTGAAAGCTA TGG |
